# Benchmarking Bioinformatic Virus Identification Tools Using Real-World Metagenomic Data across Biomes

**DOI:** 10.1101/2023.04.26.538077

**Authors:** Ling-Yi Wu, Nikolaos Pappas, Yasas Wijesekara, Gonçalo J. Piedade, Corina P.D. Brussaard, Bas E. Dutilh

## Abstract

As most viruses remain uncultivated, metagenomics is currently the main method for virus discovery. Detecting viruses in metagenomic data is not trivial. In the past few years, many bioinformatic virus identification tools have been developed for this task, making it challenging to choose the right tools, parameters, and cutoffs. As all these tools measure different biological signals, and use different algorithms and training/reference databases, it is imperative to conduct an independent benchmarking to give users objective guidance. We compared the performance of ten state-of-the-art virus identification tools in thirteen modes on eight paired viral and microbial datasets from three distinct biomes, including a new complex dataset from Antarctic coastal waters. The tools had highly variable true positive rates (0 – 68%) and false positive rates (0 – 15%). PPR-Meta best distinguished viral from microbial contigs, followed by DeepVirFinder, VirSorter2, and VIBRANT. Different tools identified different subsets of the benchmarking data and all tools, except for Sourmash, found unique viral contigs. Tools performance could be improved with adjusted parameter cutoffs, indicating that adjustment of parameter cutoffs before usage should be considered. Together, our independent benchmarking provides guidance on choices of bioinformatic virus identification tools and gives suggestions for parameter adjustments for viromics researchers.

## Introduction

Viruses of microbes (VoMs) are the most abundant life entities on Earth^1–3^, infecting many microbes at any given time^4^. The interactions between VoMs and their microbial hosts can change microbial host community composition^5^ and their physiology^6^, affecting not only the health of higher organisms (including plants, animals, and humans)^7–9^ but also the global biogeochemical processes^10,11^. Besides the direct (killing host) and indirect (e.g., recycling of limiting nutrients) ecological effects of viral lysis, VoMs can influence microbial hosts’ physiology by altering metabolic pathways through horizontal gene transfer or through the expression of metabolic genes carried within viral genomes during viral infection^6,12,13^. Understanding the complex interactions between VoMs and their microbial hosts and the ecological roles of VoMs could contribute insights on solving crucial global problems, such as infectious diseases, climate change and food crisis^14–16^. Still, improved knowledge on the genetic diversity of VoMs is warranted^17–19^. Acknowledging the genetic diversity of VoMs and their high potential for application, there is substantial interest in “mining” these viral sequences for novel anti-microbial drug candidates, enzymes for biotechnological applications, and for bioremediation^20–22^.

Originally, the discovery of most viruses depended primarily on the observation of their effect on a host. Nowadays, (meta)genomics has become a major method for virus discovery. Despite the fact that there is no universal marker gene carried by all viruses, studies of specific hallmark genes have revealed large-scale viral diversity of certain viral groups from marine and host associated biomes^23–28^. For example, Sakowski *et al*. 2014 and Wu *et al*. 2023 found great novel dsDNA bacteriophage diversity from marine samples using ribonucleotide reductase^23,28^ while Wolf *et al*. 2020, Zayed *et al*. 2022 and Edgar *et al*. 2022 expanded the known RNA viruses using RNA-dependent RNA polymerase as a marker^25–27^. Complementary to marker gene approaches, shotgun metagenomic sequencing is suitable for discovering viruses without targeted marker genes^29,30^. Through the shotgun metagenomic approach, total genetic material is extracted from environmental samples, either from an enriched viral fraction (virome) or from a fraction representing both viruses and their hosts. This is then sequenced randomly. It is common for viral reads to comprise less than 5% of metagenomic sequences^29^. Viromic datasets are derived from samples that were enriched for viruses by, e.g., size filtration, density gradient centrifugation, and/or chemical concentration^15,31,32^. Notwithstanding huge advances in virus filtration techniques^29,31,33^, it can still be challenging to distinguish viruses in metagenomic/viromic datasets due to (1) the lack of viral marker genes^23^, (2) limited availability of viral reference genomes^15^, and (3) the possible high sequence similarity between viral and microbial genomes^34^. Therefore, it has become of great interest to determine which sequences within whole microbial communities are derived from viruses. These sequences can occur as free virions, active intracellular infections, particle or host-attached virions^35^, and host-integrated or episomal viral genomes (i.e. proviruses)^29,30,36^.

In the past years, many bioinformatic virus identification tools have been developed for the task of identifying viral sequences in mixed metagenomic datasets (**Supplementary Table S1**). Some of the tools rely strongly on comparing candidate sequences to the sequences in reference databases. VirSorter^37^ uses information on the presence of viral hallmark genes, enrichment in viral-like genes, depletion in Pfam affiliated genes, enrichment in short genes and depletion in strand switching. MetaPhinder^38^ uses BLASTn and average nucleotide identity thresholds to classify viral contigs in metagenomes. Phigaro^39^, a prophage finder tool, computes the probability of each gene being localized in a prophage region based on HMMs to detect viral sequences. Sourmash^40^ uses MinHash-based sketching to create “signatures”, compressed sequence representations, that can be stored, searched, explored, and annotated. Some of the tools used machine learning techniques that also detect other genomic features based on positive and negative training sets. Seeker employs long short-term memory (LSTM) models that can identify distant dependencies within sequences, to distinguish phages from bacteria. VirFinder^41^ is a logistic regression classifier using nucleotide sequence 8-mers as features to identify viral sequences. Some of the machine learning based tools use convolutional neural networks (CNNs). DeepVirFinder^42^ and PPR-Meta^43^ use CNNs that encode long- and short-range features of viral genomes. Different from DeepVirFinder, PPR-Meta uses the BiPath algorithm that contains a “base path” and a “codon path” to learn both nucleotide and codon features. Some of the machine learning tools use hybrid approaches to combine the advantage of homology search and machine learning. VIBRANT^44^ uses viral nucleotide domain abundances in a neural network framework to classify contigs having more than four proteins. VirSorter2^45^ integrates the biological signals of VirSorter in a tree-based machine learning framework. In addition, the training/reference databases used in virus discovery are diverse, including sequence databases such as RefSeq^46^, virome datasets in specific studies^37,42,45^, and HMMs^47^ of pVOGs^39,44^. The number and diversity of virus identification tools make it challenging to choose the right tool(s), parameters, and cutoffs.

Several benchmarking studies have compared the performance of various virus identification tools^48–53^ (**Supplementary Table S2**). Most of them used simulated sequencing data or sequencing data from mock community as testing datasets. As many virus identification tools included RefSeq data in their training/reference databases, this might bias the results (more true positives). Besides, the RefSeq database contains many cultivated and isolated viral genomes while viruses in environmental samples remain under-characterized. Moreover, both the macro- and micro-diversity of viruses in natural samples is usually a lot greater than the diversity of viruses in the database, further obscuring the benchmarking results. Ho *et al*.^53^ benchmarked tools on a previously sequenced mock community containing five phage strains, which is low compared to real environmental samples. The newest benchmarking from Schackart *et al*.^52^ included two real-world metagenomic datasets but did not define a ground truth dataset.

To avoid biases in our comparison and reach the complexity level of microbial/viral communities in real metagenomic datasets, we used real-world instead of simulated or mock community metagenomic data as testing datasets. We benchmarked ten state-of-the-art bioinformatic virus identification tools on paired viral and microbial samples across three vastly distinct biomes^54^, including seawater (this study), agricultural soil^31^, and human gut^55^ (**Supplementary Table S3**). The selected viral and microbial samples were obtained through physical size fractionation, where samples went through filters with pore microbial DNA prior to virion lysis and DNA extraction in the viral fractions. Microscopic scanning or 16S amplification from viral fractions were used to confirm that there were no significant remains of microbes in the viral fraction.

We collected Illumina sequencing datasets of eight paired viral and microbial samples from each of the three biomes. First, we benchmarked the performance of tools on their default cutoffs. Second, we tested the effect of different parameters and cutoffs on the annotations. Third, we further validated our benchmarking results with extra bioinformatics tools. Our comprehensive analysis of virus identification tools in diverse biomes highlights the trade-off between specificity and sensitivity and provides valuable insights for researchers looking to identify viruses in metagenomic data.

## Methods and Data

### Construction of testing datasets from real-world metagenomic data across biomes

To investigate how the tools perform on datasets from different biomes, we selected datasets from Antarctic seawater (this study), tomato soil^31^, and human gut^55^ (**Supplementary Table S3**). We collected a total of 48 metagenome datasets, including eight paired viral and microbial datasets from each biome (**Supplementary Table S4**). The viral datasets were obtained by size fractionation through a 0.22 µm membrane followed by DNase treatment to remove free DNA, reducing the likelihood of contaminating the viruses with sequences derived from cellular organisms. To further reduce biases in the detected viruses, we selected datasets where the DNA was not amplified prior to library preparation and sequencing^56,57^. In all studies, total DNA for the microbial fraction was extracted with the PowerSoil kit. Paired-end sequencing was performed on the Illumina platform. As these microbial samples might still contain some viral sequences and the viral samples might contain some microbial sequences, any overlapping sequences between viral and microbial datasets will be removed below. The quality of each viral dataset was assessed by ViromeQC (v1.0.1)^48^, which quantifies viral enrichment in viral samples by calculating three enrichment scores based on reads mapping to the small and large subunit rRNA, and single-copy bacterial markers. Raw reads were assessed using fastp (v0.22.0)^55,58^ and MultiQC (v1.11)^59^ and assembled using metaSPAdes (v3.15.3)^60^ with k-mer sizes of 21, 33, 55, 77, 99 and 127. Contigs <1,500 bp were removed before downstream analysis using seqtk (v1.3)^61^. Raw sequencing reads were mapped back to assembled contigs with lengths at least 1,500 bp using bwa mem (v0.7.17)^62^ and the mapping statistics were summarized using samtools stats (v1.14)^63^.

Overlapping sequences between viral and microbial fractions were identified by mapping the viral to the microbial contigs using Minimap2 (v2.22)^64^ with arguments `-x ava-ont` for contigs. Contigs with Minimap hits covering at least 80% of the microbial contig length were removed from either size fraction.

We used unique contigs with lengths of at least 1,500 bp from the viral and microbial size fractions as ground truth positives and negatives, respectively. Viral contigs that were identified as viral and non-viral by the tools were regarded as true positives and false negatives, respectively. Microbial contigs that were identified as viral and non-viral were regarded as false positives and true negatives, respectively (**Figure 1**).

**Figure 1.**
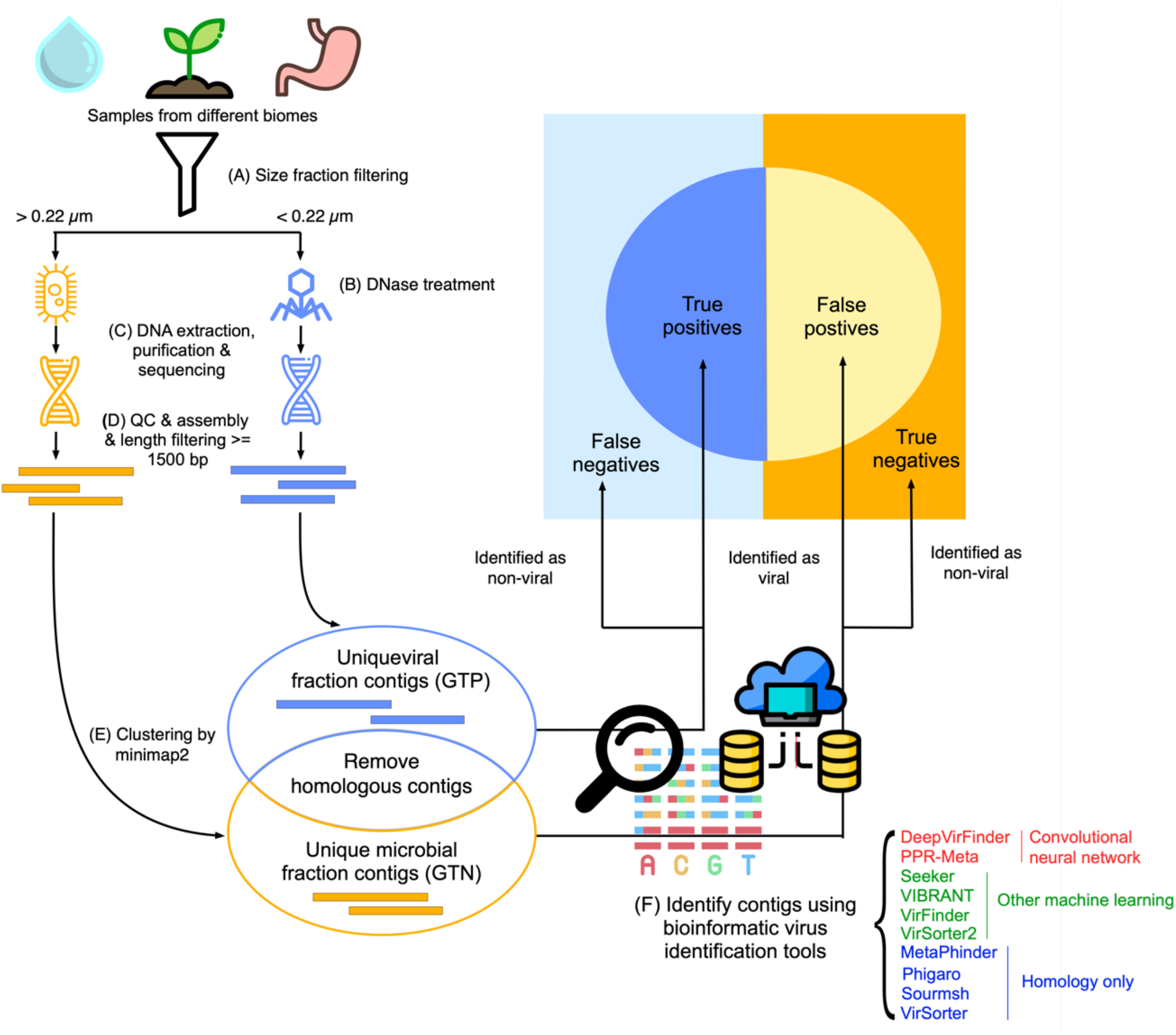
Benchmarking pipeline. (A) Samples from three different biomes were size fraction filtered through 0.22 µm filters to obtain microbial-(> 0.22 µm) and viral-enriched fractions (< 0.22 µm). (B) DNase treatment was performed in viral-enriched fractions to remove free DNA before viral lysis. (C) DNA was separately extracted, purified, and sequenced from microbial- and viral-enriched fractions to obtain viral and microbial datasets. (D) Sequenced DNA reads were quality-controlled and assembled into longer contigs. Contigs with lengths shorter than 1,500 bp were excluded from downstream analysis. (E) Homologous contigs between viral and microbial datasets were found using minimap2 and removed. Unique viral fraction contigs and unique microbial fraction contigs were used as ground truth positives and negatives, respectively. (F) Ten bioinformatic virus identification tools were applied to these datasets. Tool names were colored based on algorithms: convolutional neural network tools (red), other machine learning tools (green), and homology-only tools (blue). Viral contigs that were identified as viral and non-viral were considered as true positives and false negatives, respectively. Microbial contigs that were identified as viral and non-viral were false positives and true negatives, respectively.

### Benchmarked tools

We predicted viral contigs using the What-the-Phage pipeline (v1.0.2)^65^, which is a wrapper of eleven virus identification tools. VirNet was not run successfully. The ten remaining tools included CNN tools: DeepVirFinder v1.0^42^ and PPR-Meta v1.1^43^, other machine learning tools: Seeker with no release version^66^, VIBRANT v1.2.1 (with and without virome mode)^44^, VirFinder v1.1^41^, andVirSorter2 v2.0^45^, and homology-only tools: Metaphinder^38^ with no release version (with and without own database), Phigaro v2.2.6^39^, Sourmash v2.0.1^40^, VirSorter v1.0.6 (with and without virome mode)^37^ (**Supplementary Table S1**). The predicted classes were binarized with 0 and 1 corresponding to predicted microbial and viral origin, respectively. Our benchmarking work used contigs with lengths of at least 1,500 bp because this is the minimum length of the input sequence that can be handled by all tools. Some tools had additional requirements. For example, VIBRANT requires input sequences of at least four open reading frames. Thus, not all contigs were given a prediction result. Contigs that were not given any prediction were assigned to NAs.

### Tool performance analysis

NAs of the identification results by each tool were replaced by zero (i.e., not predicted as viral) before calculating performance measures. Performance measures, including true positive rate (TPR, also known as sensitivity, **Eq. 1**), false positive rate (FPR, **Eq. 2**), true negative rate (TNR, also known as specificity, **Eq. 3**), precision (**Eq. 4**), and F1 score (**Eq. 5**) were calculated and plotted in box plots. Receiver operation curves and UpSet plots were created using the summarized statistics.

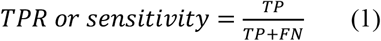

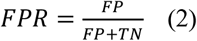

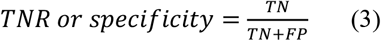

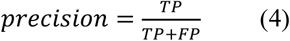

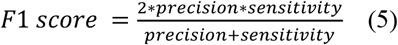

In addition, we assessed to what extent the contigs from the viral fraction represented (in-) complete genomes using CheckV (v1.0.1)^67^ and plotted the detected viral ratio in each quality rank in box plots.

### Additional contigs validation

We taxonomically classified the contigs in all testing datasets using CAT (v5.2.3)^68^ with Diamond (v2.0.9)^69^ and the CAT_database.2021-04-30 database. `CAT add_names` with the –`only_official` argument and `CAT summarise` were used to process the output. Only classifications at the superkingdom rank were used, including “Bacteria”, “Archaea”, “Eukaryota”, “Viruses”, “no support”, and “NA”. “No support” in CAT means that the ORFs in the contigs had no hit in the database. Since viruses are relatively unexplored, these contigs might be derived from novel viruses^29^. “NA” means that the ORFs had hits in the database, but these have conflicting taxonomic annotations, for example if different ORFs on a given contig map to viruses and bacteria, potentially reflecting prophages on microbial contigs. We interpreted the contigs with CAT classifications “Viruses” as potentially viral, “Bacteria”, “Archaea” and “Eukaryota”, as microbial, and “no support” and “NA” as unknown. The distributions of viral and microbial contigs in each dataset collection of contigs exclusively identified by tools or tool combinations were shown by bar plots.

To further validate the contigs in the testing datasets, we queried them for viral and microbial signals using 8,773 viral and 7,185 microbial HMMs from CheckV (v1.4)^67^. We translated the nucleotide contigs into amino acid sequences in six frames using transeq from EMBOSS (v6.6.0)^70^ and searched the sequences for the 15,958 marker HMMs using HMMsearch (v3.2.1)^47,71^. To quantify the overall viral/microbial signal in each subset of contigs, we calculated the fraction of the contig lengths that was covered by the HMM hits, converted the fraction to log ratios, and visualized the results in heatmap.

To further investigate specific sequences that were exclusively predicted by different tools, we taxonomically classified them using PhaGCN2.0 (v2.0)^72^. Gene annotation plots were created by a custom python script, after searching the translated amino acid sequences against the PHROG (v4)^73^ HMM profile database using HHsearch from the HH-suite (v.3.3.0)^74^.

All statistics were calculated and plotted using custom python and R (v4.1.0) scripts using R packages `dplyr` (v1.0.8), `tidyverse` (v1.3.1), `scales` (v0.6.0), `ggplot2` (v3.3.5), `ggbeeswarm` (v1.1.1), `ROCR` (v1.0.11)^75^, `UpSetR` (v1.4.0)^76^, `ComplexHeatmap` (v2.8.0)^77^ and `ggpattern` (v0.4.2).

### Data and scripts availability

Assembled contigs from the eight pairs of viral and microbial samples of the three biomes and other supporting files can be found in the Zenodo repository: <u>10.5281/zenodo.7692485</u>. Our full pipeline is integrated into a Snakemake (v6.5.1)^78^ workflow (**Supplementary Figure 1)**. All scripts can be found in Github repository https://github.com/MGXlab/virus_identification_tools_benchmarking.

## Results and Discussion

We benchmarked ten virus identification tools in thirteen modes using real-world metagenomic datasets. We used eight dataset pairs from each of three distinct biomes: seawater, soil, and human gut. Ground truth positives and negatives were defined as metagenomic contigs from viral (<0.22 µm) and microbial (>0.22 µm) size filters, overlapping sequences were excluded.

### Quality and composition of testing datasets from three biomes

To assess the viral enrichment and microbial contamination level of the viral datasets, we applied ViromeQC. The total enrichment score of the seawater dataset was 65 times higher than the soil dataset, and 160 times higher than the gut dataset (**Supplementary Table S5**).

Raw sequencing reads were quality-controlled and assembled into contigs. The assembly statistics, including the contig number and length distributions, are summarized in **Figure 2, Supplementary Figure 2**, and **Supplementary Table S6**. In total, seawater datasets contained the greatest number of contigs while gut datasets contained the smallest number of contigs. Seawater and gut datasets contained more microbial contigs than viral contigs while soil datasets contained more viral contigs than microbial contigs and with a greater total length. The soil datasets also had the lowest percentage of homologous contigs between the viral and microbial datasets. As overlapping viral and microbial sequences might represent active and integrated temperate viruses, respectively, we hypothesize that there might be more lytic viruses in soil than in seawater and gut, in line with previous findings that lysogeny genes are scarce in soil viromes^11,79–82^. These overlapping sequences were removed from our benchmarking analysis. Detailed information about the homologous contigs is deposited into the Zenodo repository (seawater_minimap_wtp_output.txt, soil_minimap_wtp_output.txt, and gut_minimap_wtp_output.txt).

**Figure 2.**
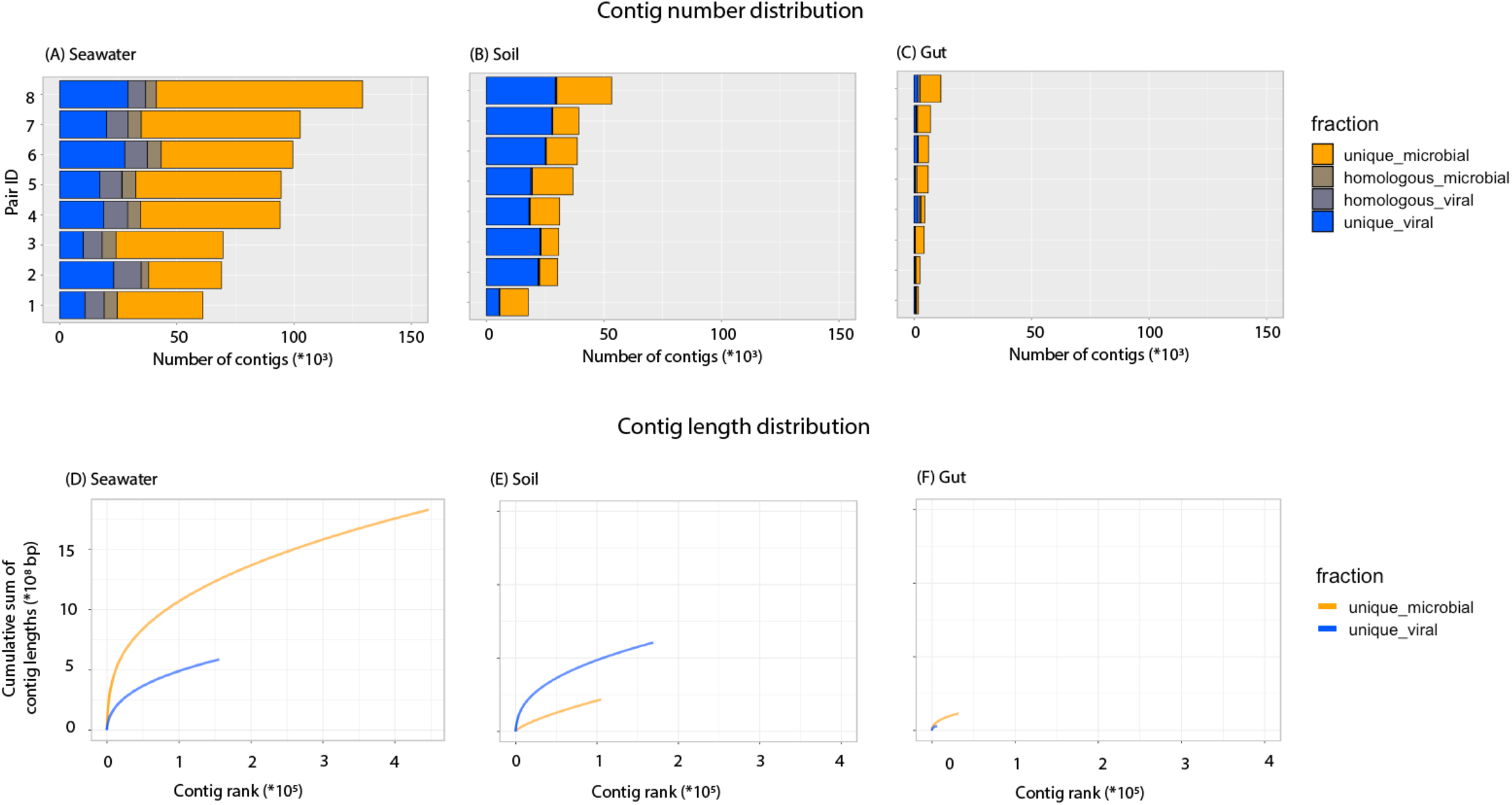
The number of contigs assembled from eight paired samples from seawater (A), soil (B), and gut (C) biomes and their cumulative lengths (D, E, F). X and Y axes of panels (A, B, C and D, E, F) were scaled to the same maximum values, so the numbers were comparable between biomes.

Assembly coverage and quality were assessed by re-mapping sequencing reads to the length-filtered assembled contigs per dataset (**Supplementary Table S7**). The percentage of properly paired reads from bwa mem mapping ranged from 11% (gut microbial dataset SRR5665119) to 94% (gut viral dataset SRR5665153) with an average of 52% (23.69%) for all datasets. Viral datasets usually had a greater percentage of properly paired reads than microbial datasets across the three biomes. This might be due to factors such as (1) the larger genomes of microbes, (2) the higher local microbial community complexity, and (3) the persistence of DNA in dead microbial matter^83,84^, which should be removed during the DNase step in the virome preparation but may persist in the microbiomes. Seawater datasets had the best assembly quality with mean percentages of properly paired reads of 79% and 45% for viral and microbial datasets, respectively. The DNA sequences of assembled scaffolds with lengths of at least 1,500 bp are deposited into the Zenodo repository (seawater_scaffolds_gt1500.fasta, soil_scaffolds_gt1500.fasta, and gut_scaffolds_gt1500.fasta). We propose that these datasets may be used in future benchmarking studies and that the results may be compared with those presented below.

### Machine learning tools outperformed homology-only tools

To compare the ability of the tools to detect viral sequences, the TPR and FPR were measured based on the percentage of contigs from the viral and microbial size fractions that were identified with default thresholds (**Figure 3** and **Supplementary Table S8**). Detailed results per contig are available in the Zenodo repository (seawater_minimap_wtp_output.txt, soil_minimap_wtp_output.txt, and gut_minimap_wtp_output.txt). All tools, except for Sourmash, detected viral contigs. Most tools ranked consistently high or low in each of the three biomes. Not many virome contigs were detected in the human gut samples by any tool. This was probably due to the contamination of microbes in the virome datasets (**Supplementary Table S5)**. Thus, optimization of experimental protocols to obtain virome from gut/fecal samples is suggested.

**Figure 3.**
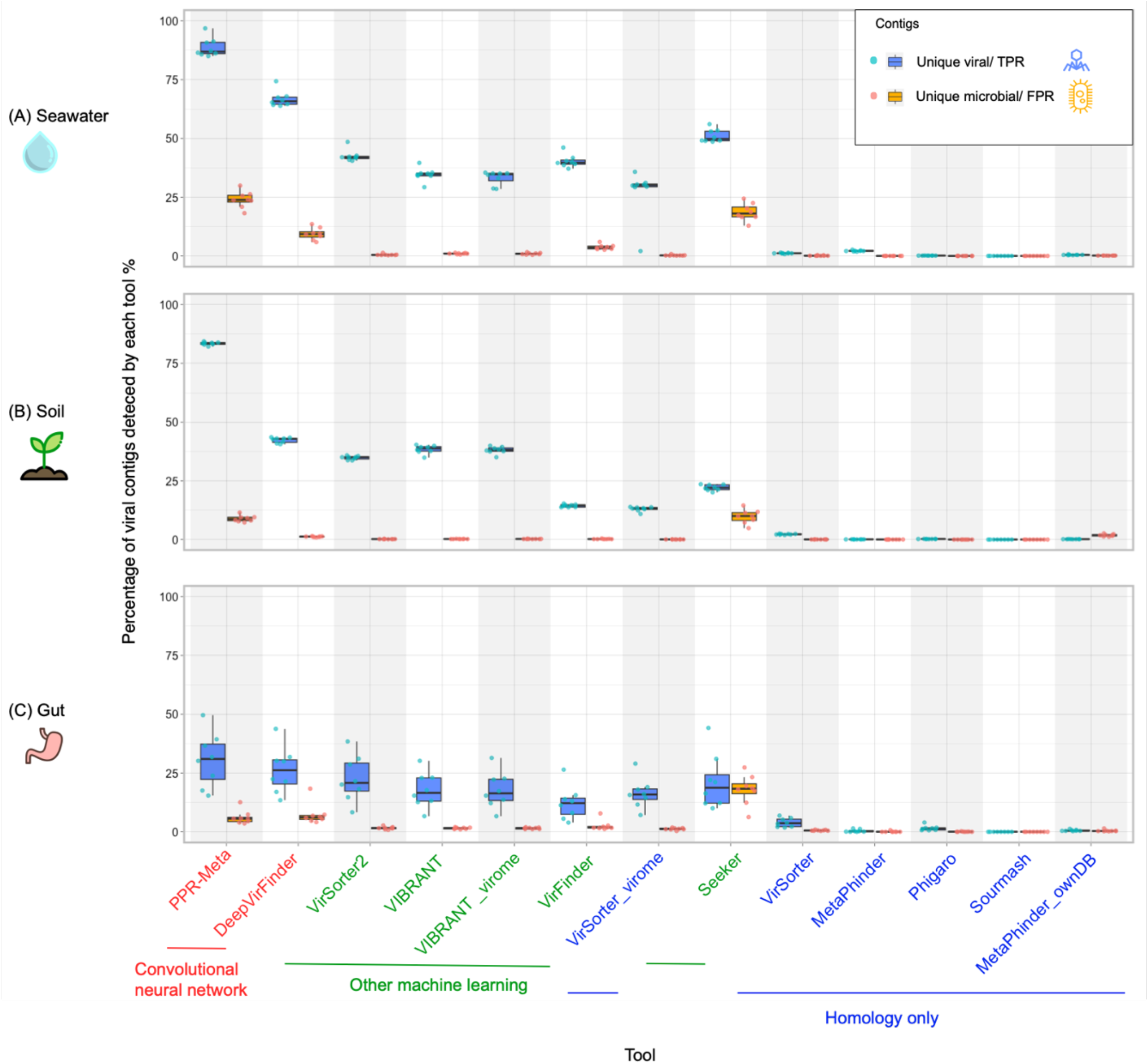
Percentage of contigs identified in the viral (TP, blue) and microbial (FP, orange) datasets, sorted by the average difference in detection rate between eight paired datasets in seawater (A), soil (B), and gut (C) biomes. Colors of the tool names as in **Figure 1**. Convolutional neural network tools outperformed other machine learning tools, while the homology-only tools performed the worst.

The choices of the algorithms and the parameters, the detected biological signals, and the compositions of the training databases all play a role in the ability of the tools to distinguish viral and microbial sequences. CNN tools outperformed other machine learning tools, while the homology-only tools performed the worst. Top performing tools, PPR-Meta (TPR/sensitivity: 68% ± 28%; FPR: 13% ± 8%) and DeepVirFinder (TPR/sensitivity: 45% ± 18%; FPR: 6% ± 5%), use convolutional neural network (CNN) algorithms that can capture short- and long-range signals on viral and microbial genomes. PPR-Meta’s CNN contains two paths -a “base path” and a “codon path” while DeepVirFinder only contains the “base path”. The “base path” is beneficial to extracting the sequence features of coding or non-coding regions while the “codon path” is specifically designed to capture the properties of coding regions^43^. The thorough detection of biological signals encoded in both DNA and codon sequences might contribute to the good performance of PPR-Meta. PPR-Meta also detected many sequences among the microbial contigs, especially from the seawater biome. This partially reflects a tradeoff between sensitivity and specificity, but it could also be that the microbial fraction contained viral elements, including integrated prophages and viruses that are actively infecting and were thus present in the microbial fraction^85^. The clustering results showed that there were many homologous contigs between the two size fractions in seawater samples (**Figure 2**), possibly indicating the prevalence of actively infecting viruses that are detected in the two fractions. Both PPR-Meta and DeepVirFinder use RefSeq viral genomes in training databases. Besides, DeepVirFinder use extra viromes from specific studies in the training database, but it did not make DeepVirFinder perform better than PPR-Meta.

VirSorter2 and VIBRANT performed similarly, identifying just under half of the viral contigs in seawater and soil metagenomes with few false positives (**Figure 2**). The two tools use similar biological signals but different algorithms. Both VirSorter2 and VIBRANT use viral domain abundance information as biological signals for identifying viral contigs. VIBRANT uses a neural network multi-layer perception classifier while VirSorter2 uses a random forest machine learning framework. Besides sequences from NCBI RefSeq, VirSorter2 includes viral sequences of giant viruses mined from public databases, novel ssDNA viruses from animals, and viral sequences from seawater biome in its training database. Focusing on the differences, VirSorter2 outperformed VIBRANT on seawater and gut biomes while VIBRANT outperformed Virsorter2 on soil biome. The inclusion of uncultivated viral sequences from human/animal tissues and seawater biomes by VirSorter2 might contribute to its better performance on the seawater and gut samples than VIBRANT (**Figure 3**).

Seeker was the only machine learning tool that did not perform well. Seeker uses much fewer parameters (∼10^2^) than PPR-Meta and DeepVirFinder (∼10^6^). The low complexity of the model, reflected in the low number of parameters might inhibit the model to capture the nuanced but important patterns in the viral sequences. Besides, like most other tools, Seeker trains on viral and bacterial sequences from NCBI RefSeq, but differently, Seeker includes many more viral than bacterial genomes in the training databases (2,232:75 and 7,375:240 viral: bacterial genomes for the first and second training databases, respectively) while the other tools include more bacterial than viral genomes (**Supplementary Table S1**). This class imbalance might lead to loss of important signals and bias the tool.

We further evaluated the TNR (specificity), precision, and F1 score of each tool (**Supplementary Figure 3** and **Supplementary Table S8**). The two CNN tools, PPR-Meta and DeepVirFinder, have higher F1 scores than the homology-only tools and other machine learning tools, mostly because of their high sensitivity, but they have lower specificity and lower precision. Homology-only tools tend to be highly specific, but they are not very sensitive and have many false negatives (undetected viral sequences). These properties should be considered when choosing a suitable tool for a specific study. For example, VirSorter2 and VIBRANT were highly specific and relatively sensitive, while PPR-Meta and DeepVirFinder were highly sensitive but less specific than the other tools. The F1 score attempts to capture both these statistics, as the harmonic mean of precision and sensitivity (**Supplementary Figure 3**). It is important to note that most machine learning tools were developed more recently than the homology only tools and therefore included more updated and complete reference databases. All tools can be expected to improve over time as newly discovered viral genomes are added to the training/reference databases. In summary, the classification algorithms, biological signals, and training/reference databases are all crucial factors in determining the performance of the tools. Both PPR-Meta and DeepVirFinder exploited CNNs, indicating the promising potential of CNN algorithms for sequence analysis.

To investigate influence of the quality of viral contigs on virus discovery, we assessed the quality of the viral contigs using CheckV (**Supplementary Figure 4**). Generally, higher quality viral sequences were easier to detect. CNN and other machine learning tools had higher sensitivity than homology-only tools on all contig quality categories. Compared to homology-only tools, machine learning tools had highly improved virus detecting ability on contigs of medium to complete quality and among which, CNN tools had highly improved virus detecting ability on contigs of low and not determined quality. PPR-Meta had high sensitivity for almost all categories of contig quality. DeepVirFinder had about half the sensitivity of PPR-Meta in all categories. VirSorter2 and VIBRANT had high sensitivity in high quality viruses (medium to complete), similar to PPR-Meta, while most low quality viruses were missed. Detailed information of the CheckV quality results of each contig is deposited into the Zenodo repository (seawater_checkv_quality_summary.tsv, soil_checkv_quality_summary.tsv, and gut_checkv_quality_summary.tsv).

### The effect of adjusting score thresholds

To further investigate the distinguishing ability of each tool, we performed Receiver Operating Characteristics (ROC) analysis. Only tools that provided a virus score were analyzed. We first grouped ROC curves by biome (**Figure 4**). For the seawater and soil biomes, PPR-Meta had the best virus discovery performance, followed by DeepVirFinder and VirSorter2. For the gut biome, VirSorter2 had the best performance, followed by DeepVirFinder and VirFinder. The high performance of PPR-Meta on soil datasets (AUC 0.95, **Supplementary Table S9**) results both from the good distinguishing ability of the tool and the quality of the testing datasets. The slightly lower performance on seawater datasets (AUC 0.90) could be a result of the tradeoff between sensitivity and specificity or of the prevalence of inactive prophages. This could have led to the detection of contigs from the microbial fraction which are counted as false positives in our benchmark (see **Figure 2** and **3**). The homology-only tool MetaPhinder performed no better than chance level based on the ROC curves.

**Figure 4.**
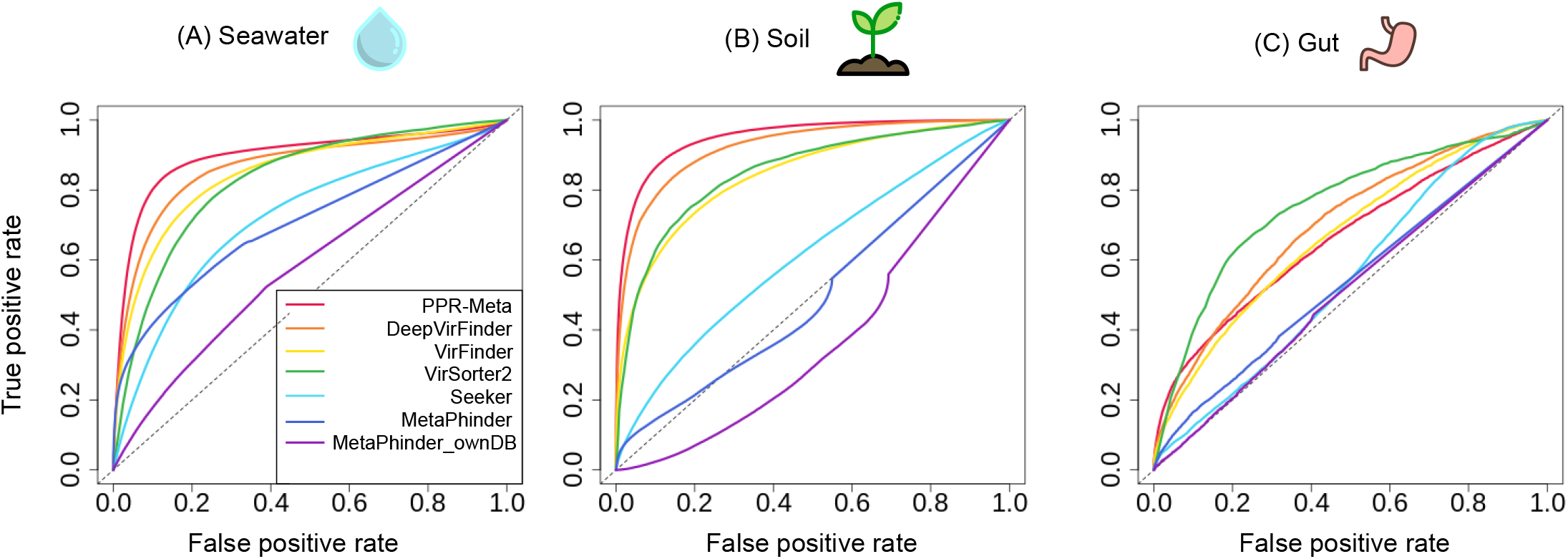
Receiver operating characteristic (ROC) curves of seven tools based on their virus identification scores in (A) seawater, (B) soil, and (C) gut biomes. The dashed diagonal line represents classification according to random chance. The area under the ROC curve (AUC) of each tool is listed in **Supplementary Table S19**.

We also grouped ROC curves by tool and colored the curves based on the cutoff score to investigate the optimal cutoff for different tools on different biomes (**Supplementary Figure 5**). The results show that adjusting virus score thresholds could enhance the virus discovery rate by some of the tools. For example, based on default thresholds, PPR-Meta and DeepVirFinder had high TPR and relatively low FPR when applied to soil datasets. Their thresholds could be decreased with a relatively low risk of false discovery so that more divergent viruses can be found. The choices of tools and thresholds depend on the goals of the study. Researchers who want to detect more novel viruses can use more sensitive tools and decrease the thresholds from the defaults, such as PPR-Meta, DeepVirFinder, and VirFinder. Researchers who have more conservative goals can use more specific tools and increase the thresholds from the defaults, such as VirSorter2.

### Agreement and disagreement between tools

To investigate the agreement and disagreement between tools, we used UpSet plots to show the intersections of identified contigs from the two size fractions of seawater (**Figure 5**), soil (**Supplementary Figure 6**), and gut (**Supplementary Figure 7**). Tools using similar algorithms clustered together and had more linkages in the UpSet matrices, indicating that the annotations of these tools tend to agree with each other. CNN tools identified the most viral contigs from the unique viral fractions and agreed with each other as they clustered on the bottom of the UpSet matrices. As expected, most unique microbial contigs were not identified as viral by any tool (large bar plots in the B panels). More consistency of predictions was seen in the viral fractions (A panels) than in the microbial fractions (B panels), indicating that tools tend to agree with each other more on viral than on microbial contigs. For seawater and soil biomes, most viral contigs were identified by at least one tool, but this was not the case for the gut biome, again suggesting that possible microbial contamination in the gut viral datasets introduced negative (microbial) sequences into the positives (viral) contig set.

**Figure 5.**
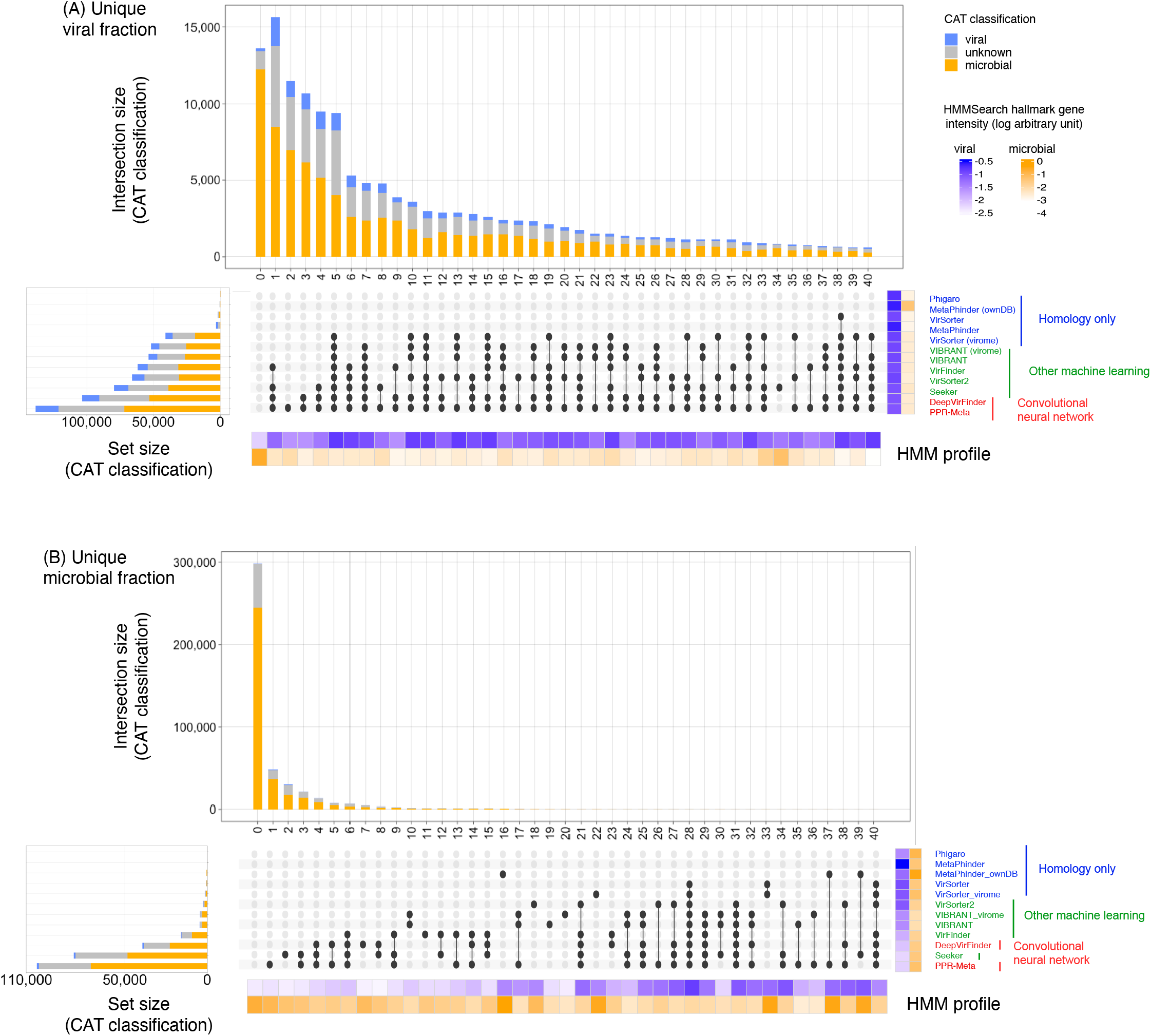
UpSet plots summarizing the overlap in predictions between tools for the unique viral (A) and unique microbial (B) contigs from the seawater samples. The total number of identified viral contigs per tool is shown in the stacked bar plots on the left. Stacked bars above of the upset plots visualize the number of viral contigs that were exclusively identified by each tool or tool combination. The left-most stacked bar shows the number of contigs that were not identified as viruses by any of the tools. The CAT classification of the contigs is indicated as colors in the bar plots: blue represents the number of contigs classified as viruses, orange represents contigs classified as “Bacteria”, “Archaea”, or “Eukaryota”, gray represents “no support” or “nan” classifications. Heatmaps below and right of the upset plots visualize the frequency of viral (blue) or microbial (orange) hallmark genes (logarithmic arbitrary units, see Methods). The intensity of hallmark gene HMM profiles was determined by dividing the length sum of all the HMM hits by the contig length. Color of the tool names as in **Figure 1**. For similar plots based on the soil and gut datasets, see **Supplementary Figures 6** and **7**, respectively.

### Characterization and annotation of predicted viral contigs

To assess the purity of the microbial and viral fractions from each biome, we further characterized the contigs using taxonomic and functional annotation tools with CAT and hmmsearch, respectively (**Figure 5** (seawater), **Supplementary Figure 6** (soil), and **7** (gut)). The stacked bars above and left of the UpSet matrices visualized the CAT classification on the contig subsets. Contigs were classified as viral or microbial by CAT, confirming an enrichment of viral sequences in the viral fraction. Summary of CAT classification on collections of contigs identified as viral by tools or tool combinations can be found in **Supplementary Table S10**. Detailed information of CAT classification results is available in the Zenodo repository (seawater_cat_taxonomy_official.txt, soil_cat_taxonomy_official.txt, and gut_cat_taxonomy_official.txt).

The heatmaps below and right of the upset plots show to what extent the contig subsets contained viral or microbial marker genes. Viral markers were found in both the viral and microbial fractions. Viral contigs that were identified by many tools contained many viral marker genes, such as column 5, 15, 23, 38, and 40 in Figure 5A. Microbial markers were mostly found in the microbial fraction, except for gut biome that contained a lot of microbial markers in the viral fraction (**Supplementary Figure 7**). A strong viral signal was also found in some microbial contigs, such as column 28 in Figure 5B which contains contigs predicted as viral by nine different tools. A summary of viral and microbial signals on contig subsets can be found in **Supplementary Table S11**. Detailed hmmsearch results can be found in the Zenodo repository (seawater_hmmsearch_domtblout.txt, soil_hmmsearch_domtblout.txt, and gut_hmmsearch_domtblout.txt).

To further investigate whether the identified contigs represent real viral sequences, the longest viral contigs that were exclusively identified by each tool in each biome were extracted, classified, and annotated. PhaGCN2 placed 6/31 of these contigs into the viral taxonomy (**Supplementary Table S12**). Although the remaining 25 contigs were not taxonomically classified, they did contain viral hallmark genes, such as genes encoding for portal proteins, head scaffolding proteins, and tail related proteins (**Figure 6, Supplementary Figure 8** and **9**), suggesting that these contigs are derived from real novel viruses. This analysis indicates that all tools have their own strengths and weakness. Some tools may not be able to predict a wide range of viruses, but almost all tools, except for Sourmash, still identified certain viral sequences that the other tools missed.

**Figure 6.**
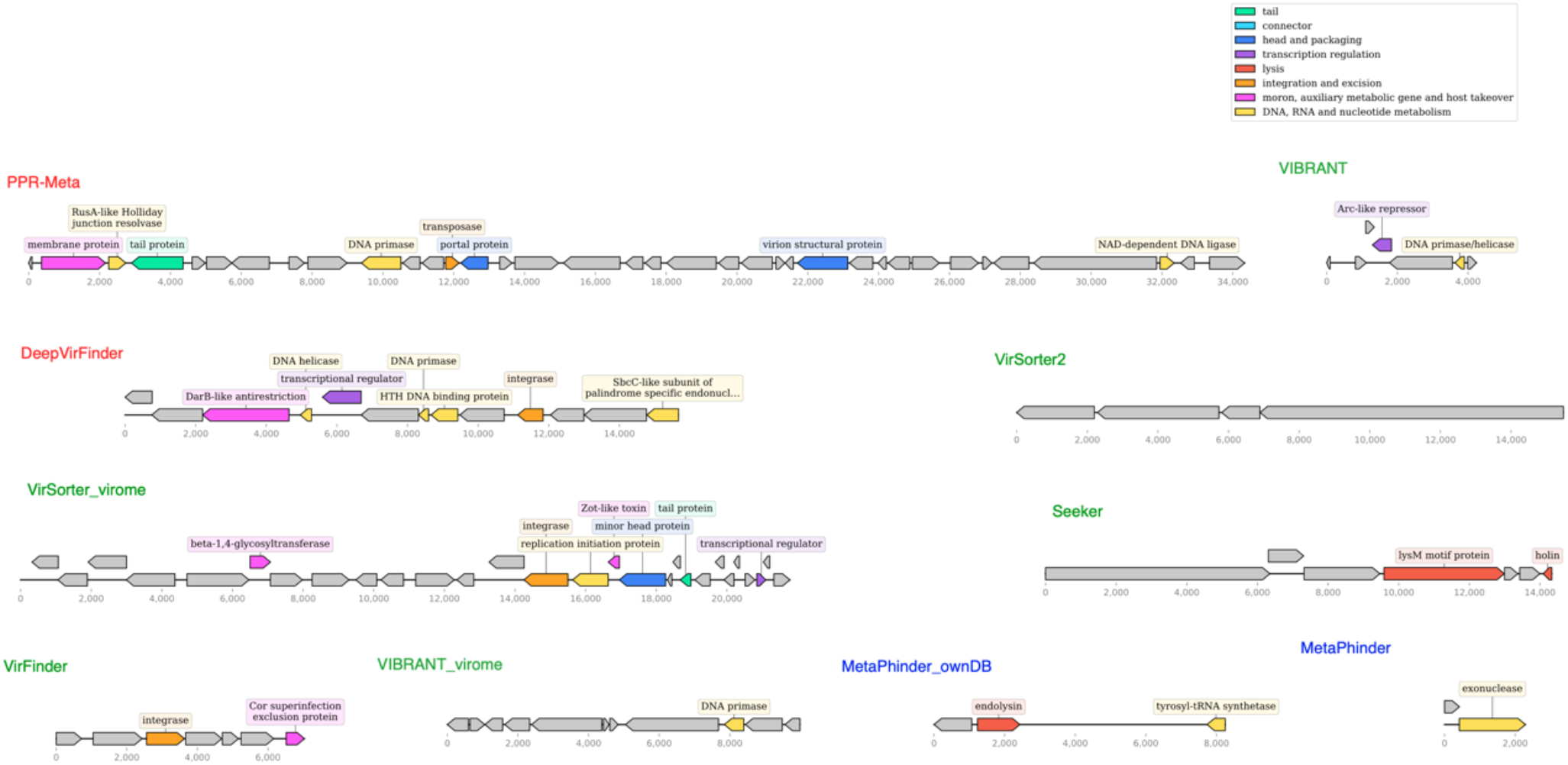
Genomic maps of the longest contigs that were exclusively identified by individual tools in the seawater virome dataset.

### Comparison to other existing benchmarking work

Several studies have compared the performance of bioinformatic virus identification tools. All studies benchmarked different sets of tools with some of them having overlaps on tested tools. One of them evaluated tools only by describing the methods of the workflows without quantification analysis^48^. Another one benchmarked tools that specialize in identifying viruses from clinical samples^51^. Four benchmarking works^49,50,52,53^ mainly used simulated viral and non-viral testing datasets that were sampled from publicly available complete viral and microbial genomes (e.g., NCBI RefSeq). A summary of the tested tools and testing datasets of each study can be found in **Supplementary Table S2**. All these studies gave insights about tool performance and which tools to choose for virus identification. Ho *et al*. 2021 found that machine learning based tools outperformed homology-only tools on the artificial dataset, which was consistent with our benchmarking results^53^. Besides a simulated dataset, they benchmarked tools on a mock community dataset consisting of five phage strains. Most tools performed significantly worse on the mock community dataset than on the artificial dataset, illustrating how artificial data sampled from RefSeq could over-estimate tools’ performance. Pratama *et al*. 2021 found that viral identification efficiency increases with fragment length, and almost all tools can correctly identify viral contigs of 10kb or longer^50^. ‘Gene content based’ tools (VirSorter) can maximize the true positive rate and minimize false positive rate at length >3 kb. K-mer based tools (DeepVirFinder) and BLAST-based tool (MetaPhinder) were good at identifying viruses from short (<3 kb) viral genome fragments. Complementary to their results, we found that the viral identification sensitivity increases with quality of contigs (**Supplementary Figure 4**). High quality contigs were better identified by machine learning tools than by homology-only tools. CNN tools were better at identifying low quality contigs than other machine learning tools. The most recent benchmarking work by Schackart *et al*. 2023 used simulated datasets as well as a real-world gut metagenome dataset^52^. Their main findings agreed with us – that homology-only tools (VirSorter, VIBRANT, and VirSorter2) demonstrate low false positive rates and robustness to eukaryotic contamination while machine learning tools have high sensitivity, allowing them to identify phages that are distinct from those in the reference databases.

For benchmarking studies that simulated testing datasets, whilst an effort was made to benchmark the tools on data that was not part of the reference/training dataset, there still might be instances of overlap. For example, Glickman *et al*. 2021 used VirSorter to perform prophage identification during testing dataset preparation, which might provide VirSorter with an advantage over other tools in prophage identification in this benchmarking work^49^. And indeed, VirSorter performed the best on identifying phages based on F1 score in this study.

The usage of newly generated metagenomic data as testing dataset in our benchmark, including a marine Antarctic dataset from an understudied region, gave us the opportunity of avoiding overlap between testing and training/reference datasets. Besides, the usage of real-world metagenomic testing datasets gives our study the perspective of metagenomic virus discovery. The main limitation of our benchmarking study is that we are not completely certain about the purity of our ground truth viral (<0.22 µm) and microbial (>0.22 µm) datasets. To address this, we carefully selected metagenomic data from the studies with appropriate standard operation procedures to separate viruses from microbes. For example, both our dataset and the publicly available datasets^31,55^ used in this benchmark performed a DNase treatment in the viral fraction before DNA extraction from virions, minimizing free microbial DNA contamination^86^. We also assessed the contamination of microbial DNA in the viral size fraction by using ViromeQC (**Supplementary Table S4**). Besides, the microbial size fraction could contain viral sequences, including integrated prophages or viruses that are attached to cells or debris. This was illustrated by the viral signals found by CAT and hmmsearch in the microbial datasets (**Figure 6, Supplementary Figure 6** and **7**). While our benchmarking work did not target prophages specifically, we minimized the influence of viral elements in the microbial size fraction by identifying and removing any homologous contigs shared between the viral and microbial fractions (**Figure 1**). We recognize that integrated prophages that are not active as free virions might still be identified as viruses in the microbial fraction (spurious false positives). We attempted to taxonomically classify some of the identified viral contigs using PhaGCN2^72^, but most of them remained unclassified. While other benchmarking studies that used simulated data had more purified data, this was at the expense of vastly reduced viral/microbial community complexity, since sequences were only sampled from a fixed set of genomes. Strain-level microdiversity, which is an important parameter in viral ecology^87^, is not represented in such simulated data. In summary, our benchmarking is complementary to the benchmarking work done by others. The usage of real-world metagenomic data from different biomes ensured the high complexity and reality of the testing datasets and avoided the problems of overlap between training and testing datasets, albeit with the compromise of not knowing the exact compositions of the testing datasets.

### Computational resources used for each tool

This study was performed on a shared high performance computing cluster. We benchmarked the computational resources used for each tool on the smallest dataset from the seawater biome (188,159 contigs, 164 MB) in a high-performance cluster of 48 Gold-6240R CPUs and 503.5 GB RAM or 48 Gold-6240R CPUs and 1510.5 GB RAM. We captured the CPU usage, physical memory usage and execution time in **Supplementary Table S13**. DeepVirFinder and VirFinder had the largest and smallest computational footprints, respectively. Two tools with good performance, PPR-Meta and VirSorter2 used intermediate computational resources. PPR-Meta is fast but relatively memory- and CPU-intensive, while VirSorter2 is relatively slow but requires less memory.

## Conclusions

This study benchmarks the performance of ten state-of-the-art bioinformatic virus identification tools using real-world metagenomic data. To evaluate how the tools perform on datasets from different biomes, we used datasets from three distinct biomes, i.e., seawater, soil, and gut. As machine learning tools outperformed homology-only tools, especially on contigs of low quality, this seems to be a very promising avenue for virus mining, and more advances are expected as this field matures. PPR-Meta, DeepVirFinder, VirSorter2, and VIBRANT performed the best among all benchmarked tools with relatively high true positive rates and relatively low false positive rates. No tool is perfect, every tool has its own strengths and weaknesses. However, it is not recommended to use union of all tools. The selection of a tool/tools may depend upon the desired application as well. PPR-Meta and DeepVirFinder were found to be sensitive but not as specific as some other tools. Thus, they can be used when detection of novel viruses is important and false positives are perhaps less of an issue. VirSorter2 and VIBRANT were more specific and precise (fewer false positives) but also less sensitive compared to PPR-Meta and DeepVirFinder. Thus, they can be applied to studies when specificity is more important than sensitivity. The adjustment of the cutoff virus scores also plays a role in the distinguishing ability of tools. If possible, in the experimental setup, we suggest using a small dataset of pairs of viral and microbial datasets for at least some samples to explore the optimal thresholds of virus detection tools using a setup as we followed in our study. For this, we have made our full Snakemake pipeline available. Besides, the experimental flow of obtaining viromes and the quality of assembled contigs would influence the discovery performance of virus identification tools.

Our comprehensive analysis of viral identification tools to assess their performance in a variety of biomes provides valuable insights to viral researchers looking to mine viral elements from novel metagenomic data across biomes. We hope that the results of this benchmarking work will provide researchers with a guide to selecting the appropriate tool and adjusting parameters for their own viral identification research.

## Acknowledgements

L.W. is funded by the Utrecht University One Health Initiative. N.P. is funded by the European Research Council (ERC) Consolidator grant 865694. Y.W. is funded by the European Union’s Horizon 2020 research and innovation program, under the Marie Skłodowska-Curie Actions Innovative Training Networks grant agreement no. 955974 (VIROINF). G.P. and C.B are funded by the Dutch Research Council NWO (grant ALWPP.2016.019). B.E.D. is funded by the European Research Council (ERC) Consolidator grant 865694: DiversiPHI, the Deutsche Forschungsgemeinschaft (DFG, German Research Foundation) under Germany’s Excellence Strategy – EXC 2051 – Project-ID 390713860, and the Alexander von Humboldt Foundation in the context of an Alexander von Humboldt-Professorship founded by German Federal Ministry of Education and Research. We acknowledge OpenAI/ChatGPT for help in proofreading our manuscript. Special thanks to Jan Kees van Amerongen and all present and past members of the Theoretical Biology and Bioinformatics group at Utrecht University and the Viral Ecology and Omics Group at Friedrich Schiller University Jena.

## Author contributions statement

L.W. designed the study, collected the data, did the analysis, built the automatic pipeline and wrote the manuscript. N.P. and Y.W. assisted with the analysis and automation of the pipeline. G.P. and C.B. contributed data. B.E.D. conceived the study and contributed to study design, data interpretation, and manuscript preparation. All authors read, edited, and approved the final manuscript.

