## supplementary figures for "Benchmarking Bioinformatic Virus Identification Tools Using Real-World Metagenomic Data across Biomes"

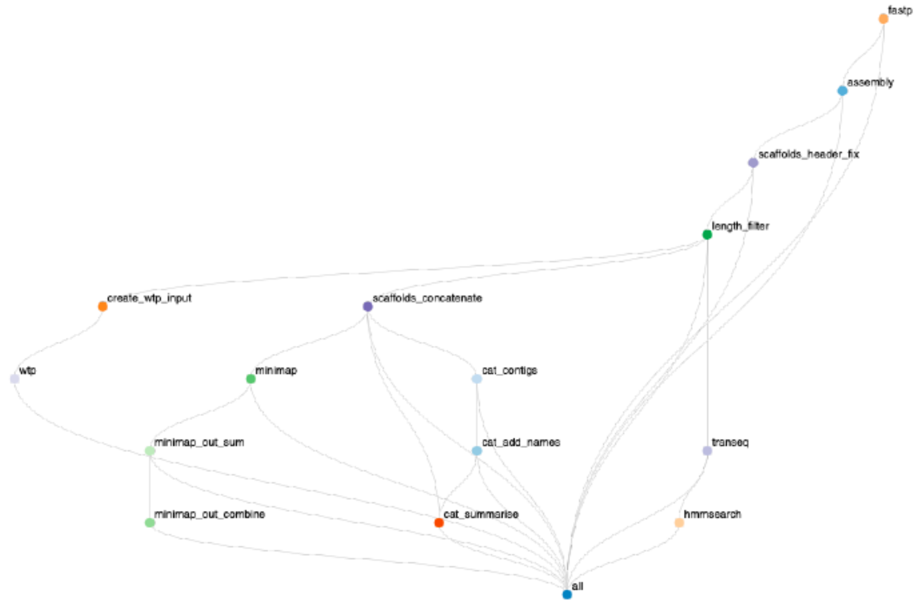


**Supplementary Figure 1.** Snakemake workflow of the pipeline to assess the quality of the raw reads, assemble the quality-control filtered reads into contigs, filter contigs based on lengths, cluster contigs from two size fractions to remove homologous contigs, run bioinformatic virus identification tools in parallel, and further validate the contigs using extra bioinformatics tools.

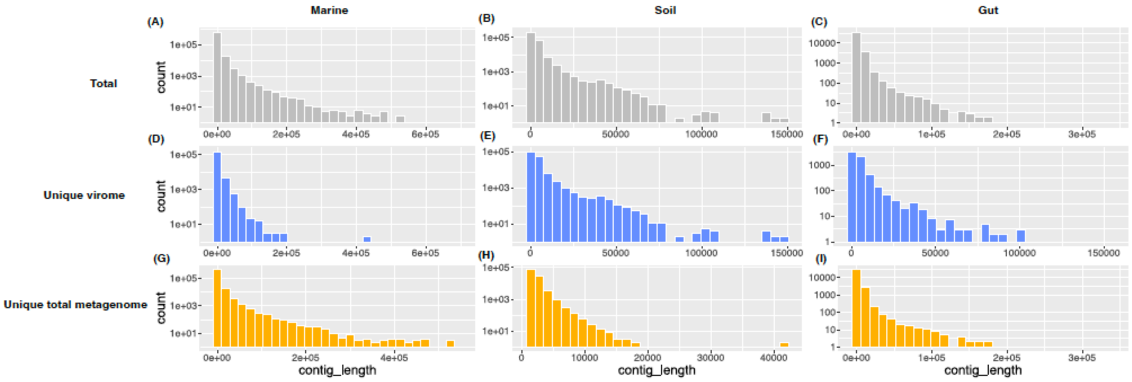


**Supplementary Figure 2.** Length distributions of all (A, B, C), unique viral (D, E, F), and unique microbial (G, H, I) contigs from seawater (A, D, G), soil (B, E, H), and gut (C, F, I) samples. Y axes are in log scales.

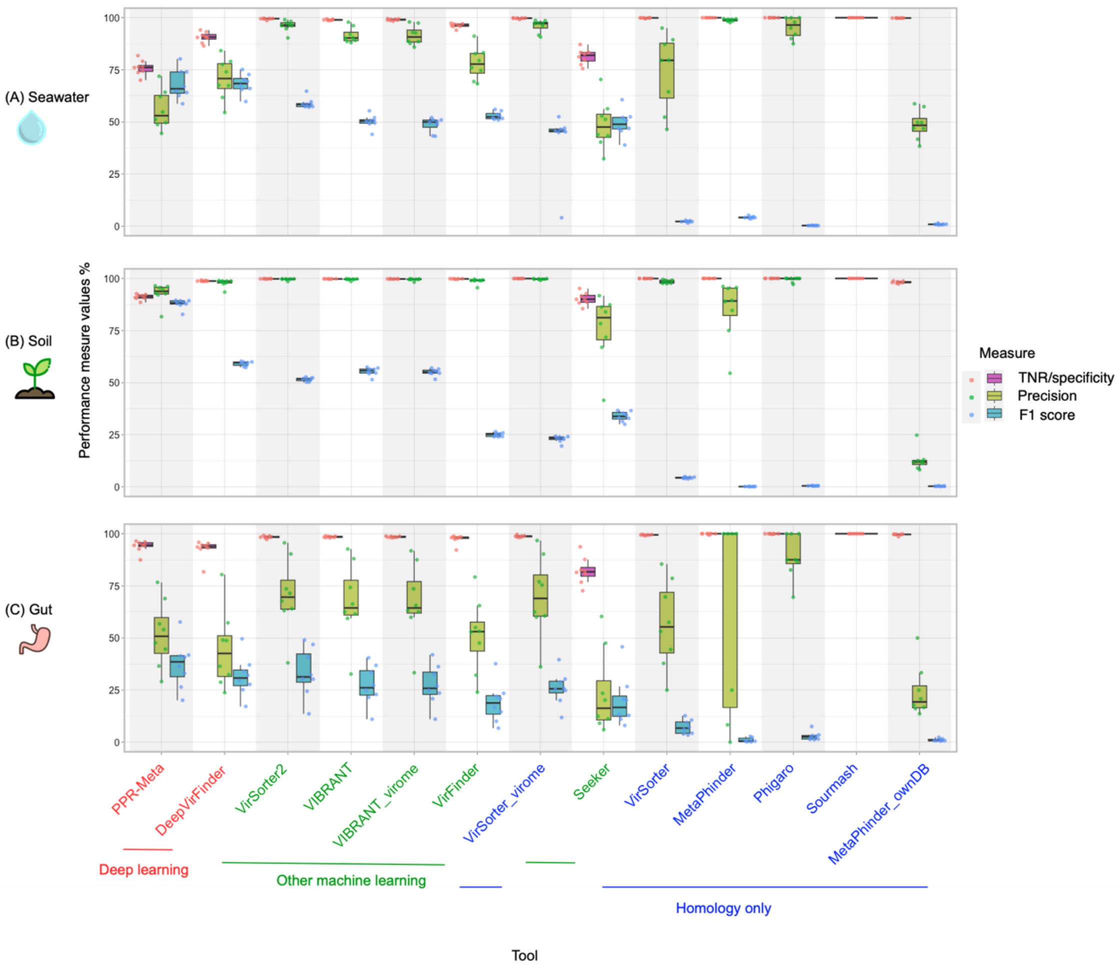


**Supplementary Figure 3.** True negative rate (TNR, also known as specificity), precision, and f1 score of each tool on samples across seawater (A), soil (B), and gut (C) biomes based on tools' default cutoffs. Sourmash did not have a precision or F1 score because it did not detect any virus. The order of the tools on the x-axis and the color of the tool names as in **Figure 3**.

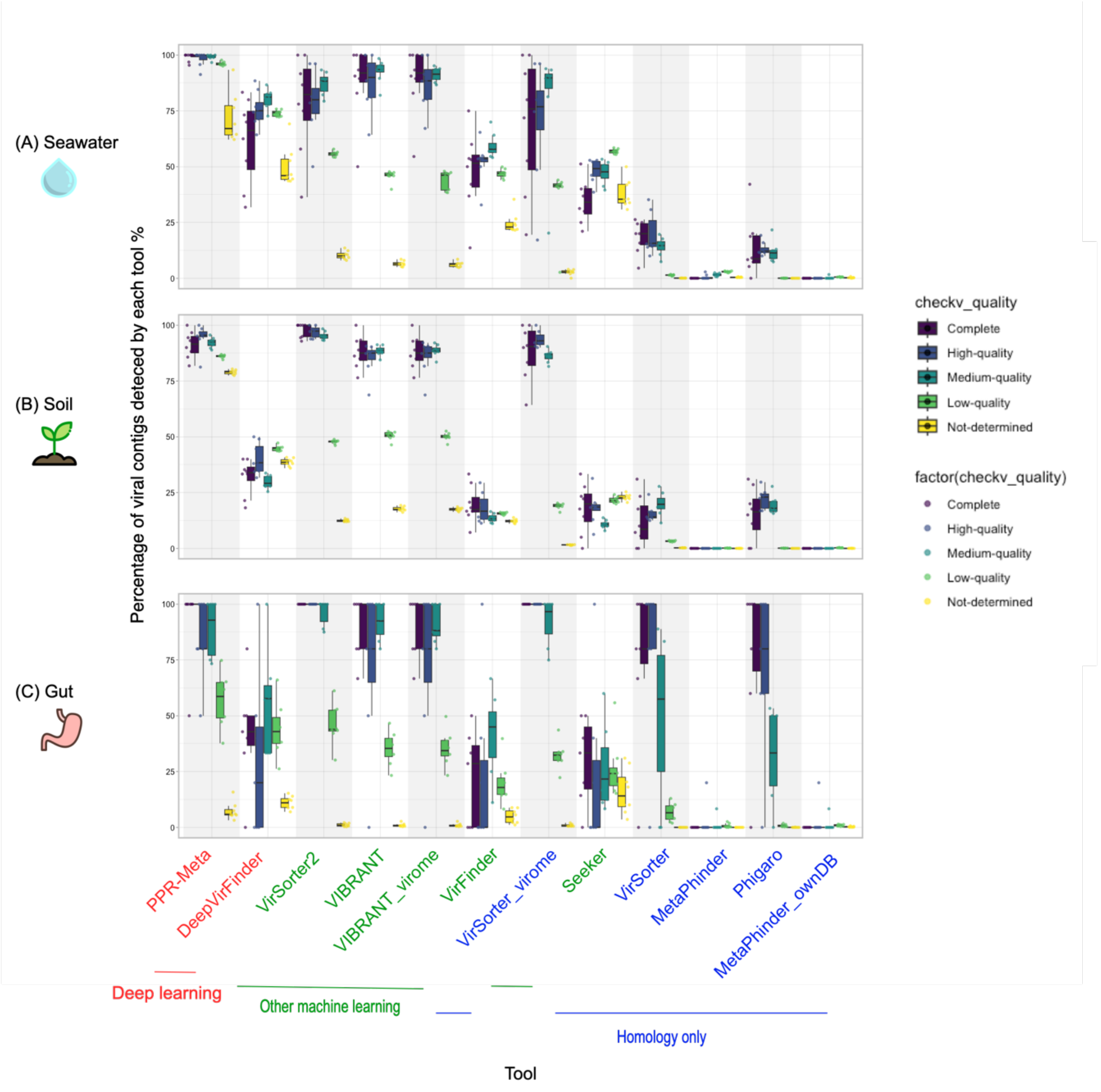


**Supplementary Figure 4.** Viral contig quality had a positive association with the viral discovery rate. Percentage of viral contigs detected by each tool from the viral fraction for viral contigs of different quality from seawater (A), soil (B), and gut (C) biomes, using default cutoffs. The order of the tools on the x-axis and the color of the tool names as in **Figure 3**.

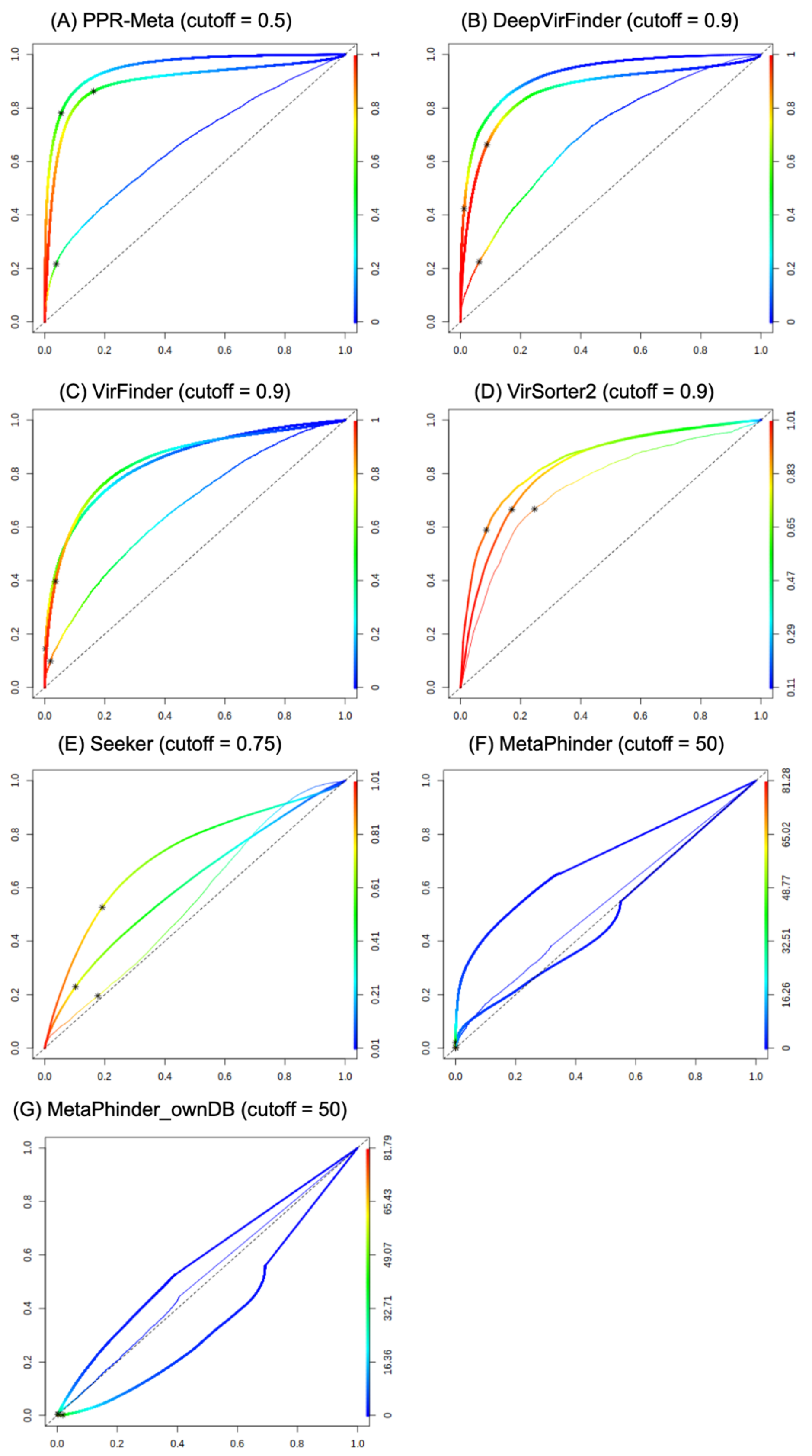


**Supplementary Figure 5.** Receiver operating characteristic (ROC) curves per tool (A) PPR-Meta, (B) DeepVirFinder, (C) VirFinder, (D) VirSorter2, (E) Seeker, (F) MetaPhinder, and (G) MetaPhinder with own database. The curves from outside to inside were for soil, seawater, and gut biome, respectively. Asterisks were the default cutoffs (values shown in the header of each panel) of tools. Color scheme showed all the possible cutoffs of each tool. The area under the ROC curve (AUC) of each tool is listed in **Supplementary Table S9**.

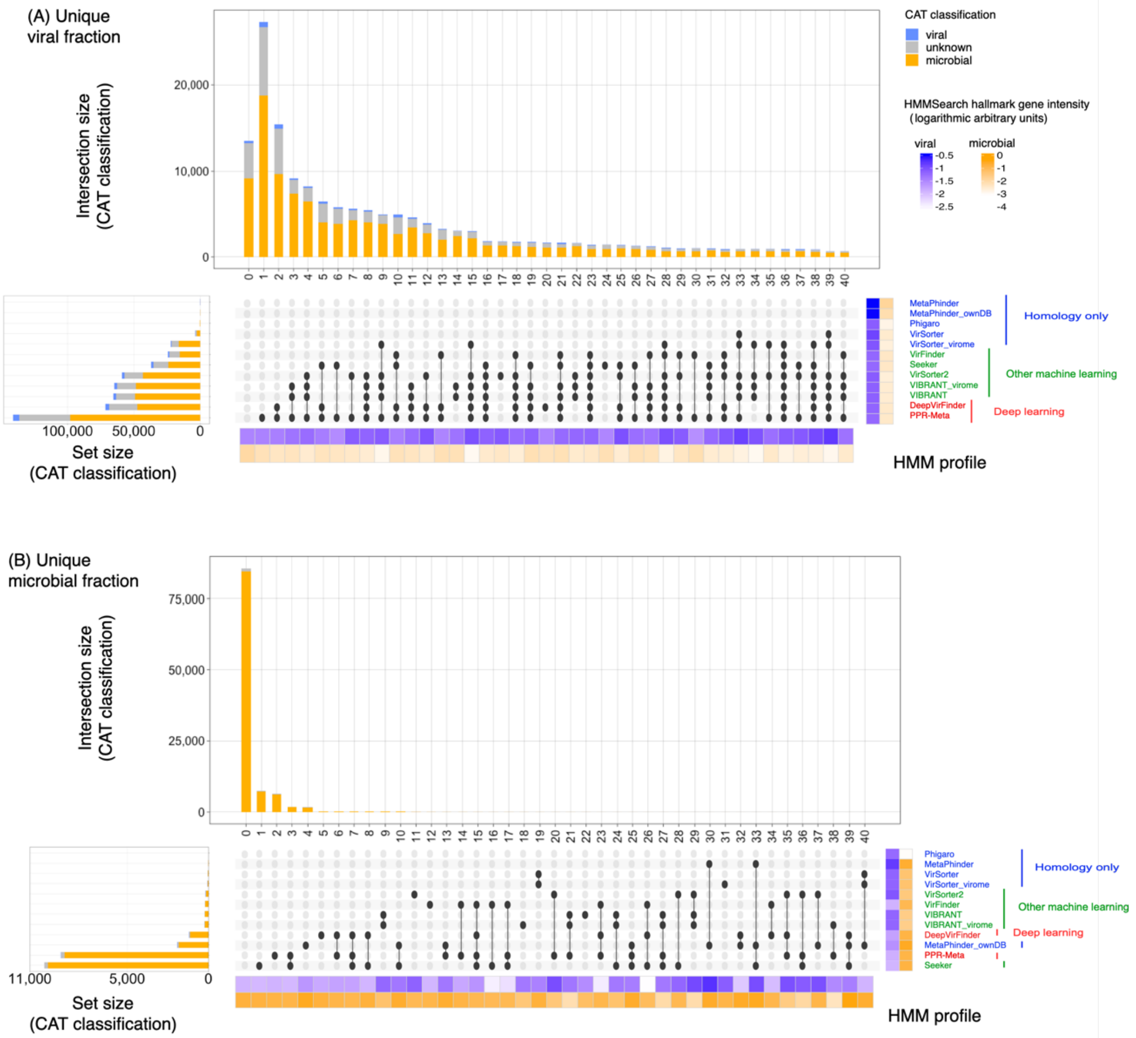


**Supplementary Figure 6.** UpSet plots summarizing the overlap in predictions between tools for the unique viral (A) and unique microbial (B) contigs from the soil samples. The total number of identified viral contigs per tool is shown in the stacked bar plots on the left. Stacked bars above of the upset plots visualize the number of viral contigs that were exclusively identified by each tool or tool combination. The left-most stacked bar shows the number of contigs that were not identified as viruses by any of the tools. The CAT classification of the contigs is indicated as colors in the bar plots: blue represents the number of contigs classified as viruses, orange represents contigs classified as “Bacteria”, “Archaea”, or “Eukaryota”, gray represents “no support” or “nan” classifications. Heatmaps below and right of the upset plots visualize the frequency of viral (blue) or microbial (orange) hallmark genes (logarithmic arbitrary units, see Methods). The intensity of hallmark gene HMM profiles was determined by dividing the length sum of all the HMM hits by the contig length. Color of the tool names as in **Figure 1**.

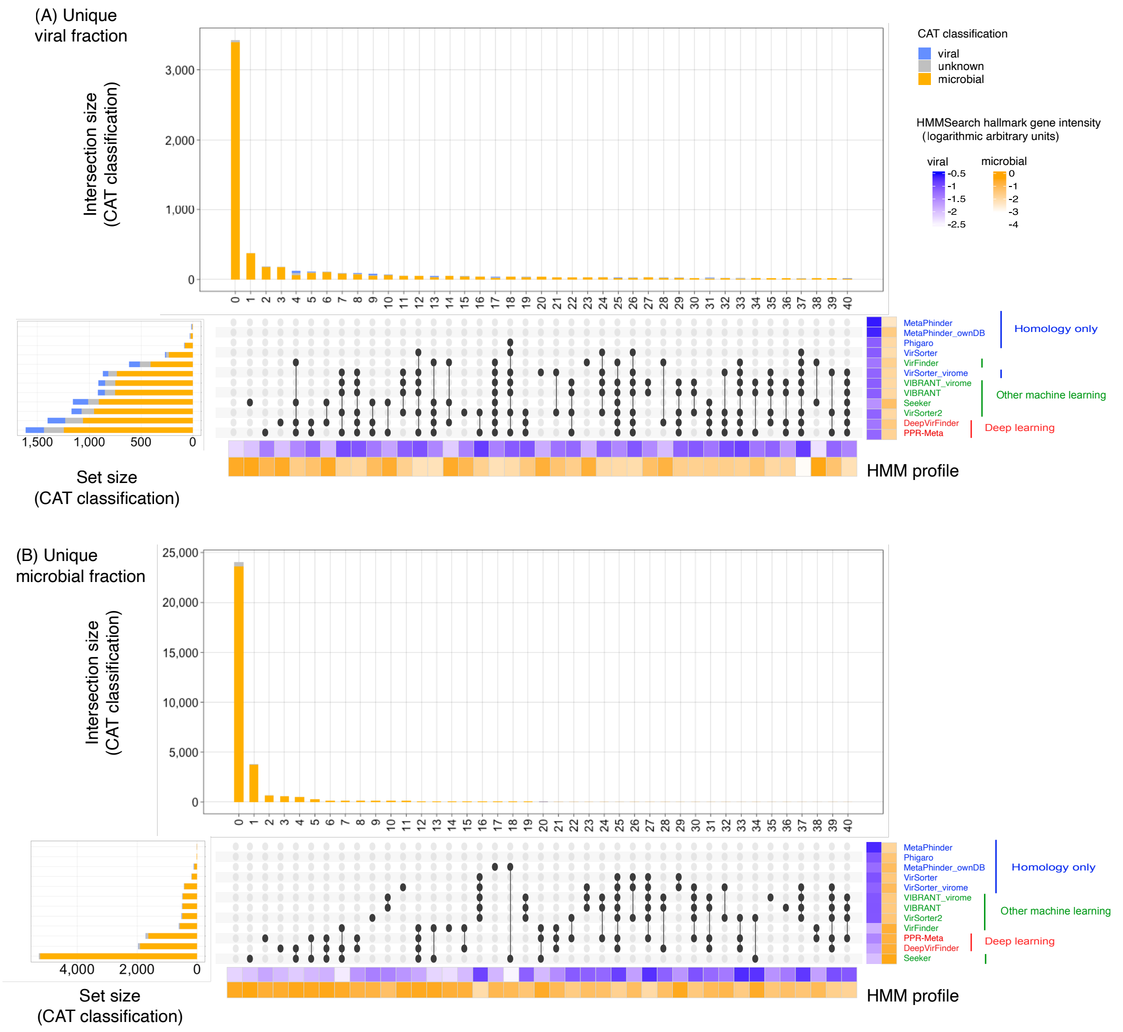


**Supplementary Figure 7.** UpSet plots summarizing the overlap in predictions between tools for the unique viral (A) and unique microbial (B) contigs from the gut samples. The total number of identified viral contigs per tool is shown in the stacked bar plots on the left. Stacked bars above of the upset plots visualize the number of viral contigs that were exclusively identified by each tool or tool combination. The left-most stacked bar shows the number of contigs that were not identified as viruses by any of the tools. The CAT classification of the contigs is indicated as colors in the bar plots: blue represents the number of contigs classified as viruses, orange represents contigs classified as “Bacteria”, “Archaea”, or “Eukaryota”, gray represents “no support” or “nan” classifications. Heatmaps below and right of the upset plots visualize the frequency of viral (blue) or microbial (orange) hallmark genes (logarithmic arbitrary units, see Methods). The intensity of hallmark gene HMM profiles was determined by dividing the length sum of all the HMM hits by the contig length. Color of the tool names as in **Figure 1**.

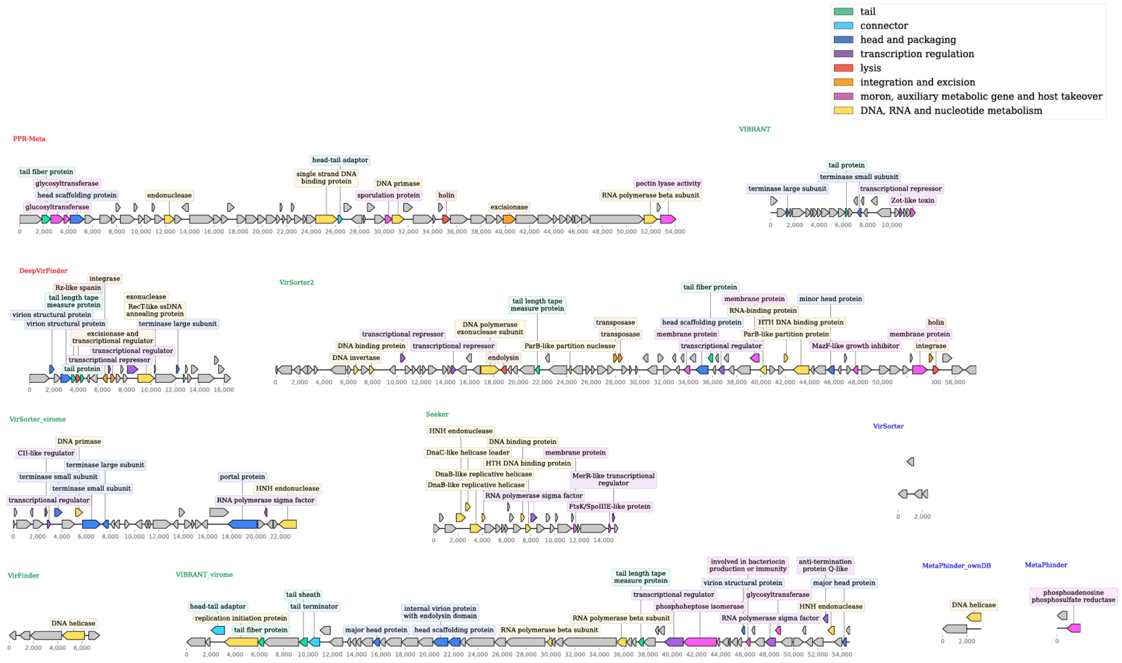


**Supplementary Figure 8.** Genomic maps of the longest contigs that were exclusively identified by individual tools in the soil virome dataset.

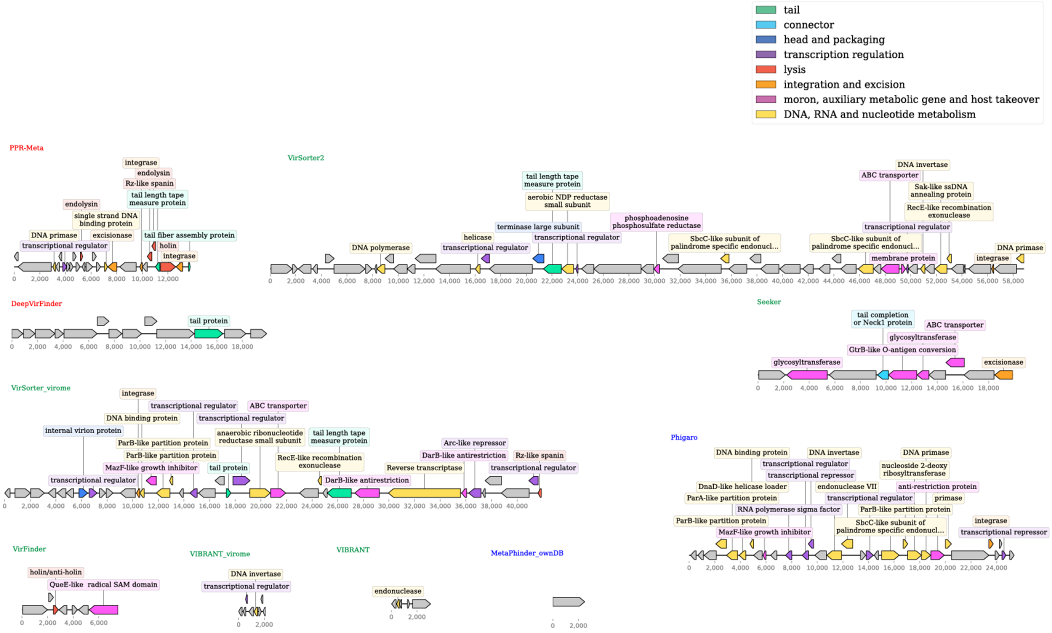


**Supplementary Figure 9.** Genomic maps of the longest contigs that were exclusively identified by individual tools in the gut virome dataset.
